# Pharmacological manipulations of judgement bias: a systematic review and meta-analysis

**DOI:** 10.1101/612382

**Authors:** Vikki Neville, Shinichi Nakagawa, Josefina Zidar, Elizabeth S. Paul, Malgorzata Lagisz, Melissa Bateson, Hanne Løvlie, Michael Mendl

## Abstract

Validated measures of animal affect are crucial to research spanning a number of disciplines including neuroscience, psychopharmacology, and animal welfare science. Judgement bias, which assesses decision-making under ambiguity, is a promising measure of animal affect. One way of validating this measure is to induce affective states using pharmacological manipulations and determine whether the predicted judgement biases are observed. Here, we conducted a systematic review and meta-analysis using data from 19 published research articles that use this approach from which 440 effect sizes were extracted. The results of the meta-analysis suggest that pharmacological manipulations overall altered judgement bias as predicted. However, there were several moderating factors including the neurobiological target of the drug, whether the drug was hypothesised to induce a relatively positive or negative affective state, dosage, and the presented cue. This may partially reflect interference from adverse effects of the drug, such as sedation. Thus, while judgement bias can be used to measure pharmacologically-induced affective states, potential adverse effects of the drug should be considered when interpreting results.

## 1. Introduction

Measurement of affective state, which is defined as comprising both shortterm emotions and longer-term moods [1], is important to a number of disciplines including psychopharmacology, neuroscience, and animal welfare science, as well as being of societal interest. Mood disorders, for example, are a significant global concern; it is estimated that 780,000 people died by suicide in 2015, with one death every 40 seconds on average [2]. Major depressive disorder is ranked as the largest single contributor to global disability, and anxiety disorders are ranked sixth [2]. The development of pharmacological treatments for mood disorders has been largely dependent on the use empirical studies using non-human animals. Reliable and validated measures of affective state in non-human animals are therefore crucial to understanding the neurobiological aetiology of these disorders and to the further development of treatments. In particular, measures should have both predictive validity (i.e. the extent to which the measure is altered in the predicted direction by drugs which alter human affect) and construct validity (i.e. the extent to which they measure precisely what they claim to measure) [3]. Predictive validity is typically regarded as the gold-standard for validating novel behavioural measures of affective state.

Numerous behavioural assays have been developed to assess animal affect. The most common of these include the forced swim test, and its derivative, the tail suspension test, which are considered to measure helplessness [4, 5, 6, 7]; the sucrose preference test which is considered to measure hedonic capacity [8, 9]; and the elevated plus maze considered to assess the relative of value of exploration to safety [10, 11]. Overall, there is good evidence to suggest that these assays have predictive validity with a broad range of antidepressant or anxiolytics drugs resulting in changes in the predicted direction. Although, antidepressants that are used to treat generalised anxiety disorder in humans do not consistently alter rat behaviour in the elevated plus maze [12, 13, 14]. However, the construct validity of these assays has been disputed. For example, it has been argued that the forced swim test and tail suspension test reflect a learnt response rather than helplessness [15, 16, 17, 18, 19]. Similarly, research has shown that humans with depression show no reduction in their preference for sucrose over water [20, 21] and that body weight may be a strong confounding factor in the sucrose consumption test [22]. The outlined deficiencies in currently used assays means that there is a clear need for improved methods to measure affective state in non-human animals that have both construct and predictive validity.

The judgement bias task (sometimes referred to as the cognitive bias task) provides an alternative means to examine affect in non-human animals and has been used widely in the field of animal welfare science since its conception by Harding et al. (2004) [23, 24, 25]. Rather than specifically measuring anxiety or depression, the task purports to measure affective valence; the relative pleasantness or unpleasantness of the current state of the individual. To achieve this, the judgement bias task examines decision-making under ambiguity. Individuals are first trained to associate the presentation of one reference cue (e.g. a high frequency tone) with a reward and presentation of another reference cue (e.g. a lower frequency tone) with a lower reward or punisher. Once training is complete, individuals are presented with one or a few untrained probe cues that are intermediate between the reference cues (e.g. medium frequency tones). Whether an individual responds to these ambiguous cues as though they signal the more positive or less positive outcome is considered to be indicative of their expectation of rewards or punishers and hence affective state. Less risk-averse individuals (often deemed ‘more optimistic’ or ‘less pessimistic’) are considered to be in a relatively more positive affective state.

The task is based on the empirical finding that humans experiencing anxiety and depression have a greater expectation of punishing events or reduced expectation of rewarding events than clinically healthy humans [26, 27, 28]. Moreover, back-translation of the task to human subjects has demonstrated a correlation between judgement bias and measures of subjectively-experienced affect, such as the State-Trait Anxiety Inventory (STAI), Visual Analogue Scale for Anxiety (VAS-A), and negative affect dimension of the Positive and Negative Affect Schedule (PANAS) [29, 30, 31]. The task therefore appears to have strong construct validity.

Research has been conducted to assess how judgement bias is influenced by affect-altering drugs in non-human animals. Synthesis of these studies would allow conclusions to be drawn about the ability of the judgement bias task to measure pharmacologically induced positive (relatively more pleasant) or negative (relatively less pleasant) affective states. To this end, we conducted a systematic review and meta-analysis to assess whether pharmacological manipulations alter judgement bias and hence assess the predictive validity of the task. In addition to assessing whether there was an overall effect, we investigated whether the relationship between affect-altering drugs and judgement bias was moderated by factors relating to the drug and administration of the drug, such as the duration and timing of administration, dosage, and neurobiological target of the drug (see Box 1). The potential moderating effects of several task-related factors, such as the presented cue, species used, sex, reinforcement type, response type, and the outcome measure, were also investigated. While we predicted that the effects of judgement bias would be largest at the ambiguous cues and would depend on dosage, we did not predict that the other moderators would influence the effect of the pharmacological manipulations on judgement bias.

**Box 1: Neurobiological targets of affect-altering drugs:**

*Adrenergic system*: Epinephrine and norepinephrine are both hormones and neurotransmitters that bind to adrenergic receptors. The adrenergic system is involved in the early stages of a stress response [32]. Brains of depressed patients have reduced levels of norepinephrine and antidepressant drugs such as reboxetine selectively-target the adrenergic system [33, 34].

*Dopaminergic system*: Dopamine is a neurotransmitter and neuromodulator that can have both inhibitory and excitatory effects on target dopamine neurons. Dopaminergic-system dysregulation is associated with depression [35]. Antidepressants targeting dopamine (although non-specifically) such as monoamine oxidase inhibitors (MAOIs) are available but limited in clinical usage [35].

*Gamma-Aminobutyric acid (GABA) system*: The neurotransmittor GABA, which binds to GABA receptors, is the major inhibitory neurotransmitter in the brain [36]. Reduced GABA levels are associated with panic disorder [37, 36]. A number of commercially available treatments for anxiety disorders, such as barbituates, benzodiazepines, and gabapentins, are purported to work by enhancing GABA function.

*Glucocorticoid system*: Glucocorticoids are a class of steroid hormones that bind to glucocorticoid receptors. The system is involved in the later stages of a stress response, altering cognitive functioning, such as attention and memory, following an acute stressor [32].

*Glutaminergic system*: Glutamate is the brains major excitatory neurotransmitter and targets glutaminergic receptors that include NMDA, AMPA, and kainite. Several recreational dissociative drugs target the glutaminergic system specifically, such as ketamine and phencyclidine (PCP) [38]. *Opioid system*: Opioid receptors are targeted by a number of neuropeptides including endorphins and nociceptin. The opioid system plays a key role in pain modulation, and mediates the euphoric and analgesic effects of a number of recreational and clinical drugs such as morphine and heroin [39].

*Oxytocin system*: Oxytocin is a hormone and neuropeptide that targets the oxytocin receptor. The oxytocin system has been implicated in depression; low oxytocin levels have been observed in depressed patients [40].

*Serotoninergic system*: Serotonin is a primarily inhibitory neurotransmitter that binds to serotonergic receptors [41]. There is a wealth of evidence indicating a link between low levels of serotonin and depression [41]. Antidepressant drugs that target the serotonergic system, such as citalopram and fluoxetine, are commonly prescribed [42].

## 2. Methods

### 2.1. Literature search

This study followed the Preferred Reporting Items for Systematic reviews and Meta-Analyses (PRISMA) statement (see Fig. 1)[43]. In addition to including all research articles that had been excluded from the meta-analysis by Nakagawa et al (*in prep.*) for using pharmacological manipulations of affective state, a literature search was conducted on the 13th November 2017 using Scopus and Web of Science to identify more recent research papers. This search excluded results prior to 2016 as the search by Nakagawa et al (*in prep.*), which used the same search-terms, had already identified pre-2016 research articles.

**Figure 1:**
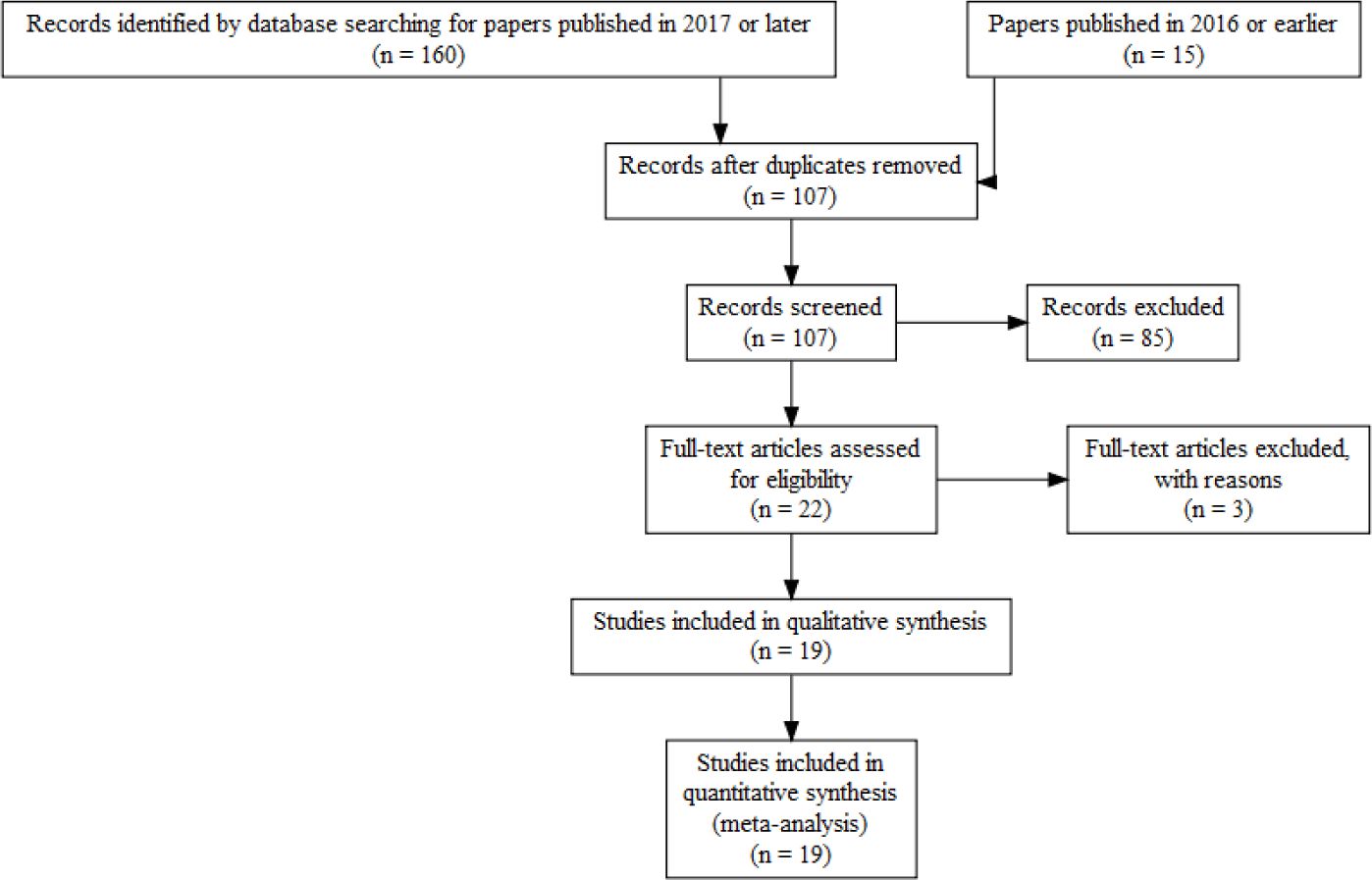
PRISMA Flow Diagram illustrating the number (n) of articles included at each stage of the literature review.

The search-terms used in Scopus and Web of Science were: (”Cognitive bias*” OR “judgment bias*” OR “judgement bias*” OR “Cognitive affective bias*”) AND (”pessimis*” OR “optimis*” OR “valence” OR “mood*” OR “emotion*” OR “affective state*” OR “emotional state*” OR “ambig*”) AND (”animal*” OR “animal welfare”). These searches were conducted within the titles, abstracts, and keywords of research articles.

### 2.2 Inclusion and exclusion criteria

The literature search identified 160 research articles; 94 from Scopus and 66 from Web of Science. After removal of duplicate articles, and inclusion of the articles using pharmacological manipulations of affect that were excluded by Nakagawa et al (*in prep.*) in their meta-analysis, there were 107 articles to be screened. These articles were first screened solely by their abstract. During this abstract-based selection, articles were selected for tentative inclusion in the meta-analysis if they were deemed to be an empirical study which compared judgement bias between at least one control group and at least one treatment group to whom an affect-altering drug had been administered. Additionally, to be included, these studies had to be conducted on vertebrate, non-human animals. An affect-altering drug was any substance that was considered to have antidepressant, depressant, anxiolytic, or anxiogenic effects. Twenty-two articles met these inclusion criteria.

In the full-text screening, articles were selected on the basis that they had used a variant of Harding *et al*’s (2014) cognitive judgement bias task to compare judgement bias between a group of individuals to whom a vehicle substance had been administered and at least one treatment group who had been given a drug to induce a positive or negative shift in their affective state [23]. To be included, the outcome measure had to be either latency to approach the cue (e.g. a location in a test arena) on each trial, where approaching the presented cue had been associated with reward and hence shorter latencies would be interpreted as greater ‘optimism’, or proportion of positive responses to each presented cue where a greater proportion would be interpreted as greater ‘optimism’, or an outcome measure that could be converted into either form. For example, if the article reported the proportion of negative responses to each presented cue or reported the percentage of positive responses to each presented cue, the extracted data would be subtracted from one or divided by 100 respectively. All included articles either reported the proportion or latency, but not both, and hence only one of these measures was extracted for each article. Two articles were excluded at the full-text selection stage following retraction by their authors. In addition to these two exclusions, one author did not respond to our request for data and hence data from their article could not be included in the meta-analysis. A total of 19 articles were included in the meta-analysis.

### 2.3. Data extraction

We extracted the mean and standard deviation of either the latency to approach the presented cue, or proportion of positive responses to the presented cue, as well as the sample size, for both the treatment and control group for each cue from each article (JZ and VN extracted the data which were checked by VN and SN). Data in a graphical format were extracted using GraphClick 3.0.3 (http://www.arizona-software.ch/graphclick/) or WebPlotDigitizer 4.1 (http://automeris.io/WebPlotDigitizer). If multiple drug doses had been used, these variables were extracted for each dosage, and similarly, if there were test sessions that varied the duration between administration and testing or number of days of chronic drug administration, these variables were extracted for each test session. Data collected from both a vehicle and treatment group prior to drug administration were not included as these were not informative. In cases where data could not be extracted from the article or supplementary material, the data were requested from the corresponding author.

The extracted treatment and control group data were categorised as expected to induce either a more or less positively-valenced affective state. If an anxiogenic or depressant substance had been administered, as determined by the hypotheses stated in the published article alongside the information outlined in Box 1, the treatment group was categorised as the less positive group, and the vehicle group was categorised as the more positive group. If an anxiolytic or antidepressant drug had been administered, which was also determined by the hypotheses stated in the article alongside the information outlined in Box 1, the treatment group was categorised as the more positive group and the vehicle group categorised as the less positive group. If no hypotheses were stated in the article, this categorisation was based on the description and pharmacodynamics of the substance as outlined on the DrugBank database [44] in addition to the information presented in Box 1. Where multiple doses had been administered, higher doses of anxiolytic or anxiogenic drugs were categorised as more positive whereas higher doses of anxiogenic or depressant drugs were categorised as less positive.

Information about the article and authors, drug and drug administration, and methodology were also extracted (Table 1). These included; the article title, institute or university at which the research was conducted, the name of the drug, the dosing duration (chronic - where drugs were administered repeatedly, acute - where the drug was administered immediately before testing, or chronic wash-out the period after drug administration had stopped following chronic administration), the time between administration and testing (acute studies only), the number of days since the first dose (chronic studies only), the dosage (in mg/kg), the neurobiological target of the drug, the manipulation type (positive or negative affect induction), the species tested, and the outcome variable used (latency or proportion), cue (positive reference cue, midpoint probe cue, negative reference cue, and where included the near negative probe cue and near positive probe cue), sex of the experimental subjects (all male, all female, or both male and female), reinforcement type which was the type of reinforcement used for the reference cue (reward-punishment - where the positive reference cue was rewarded and negative reference cue punished; reward-null - where the positive reference cue was rewarded and negative reference cue was not rewarded; or reward-reward - where the positive reference cue was rewarded with a high reward and negative reference cue was reward with a low reward), response type which reflected whether both or only one of the reference cues required an approach response (go/no-go - where the positive reference cue required an active response and the negative reference cue required no response, or go/go - where both reference cues required an active response), and cue type (reference or probe). To ensure that dosage was comparable between substances and species, each drug dose within a species was standardized by dividing the dosage (in mg/kg) by the standard deviation of all doses administered within each drug for each species.

**Table 1:**
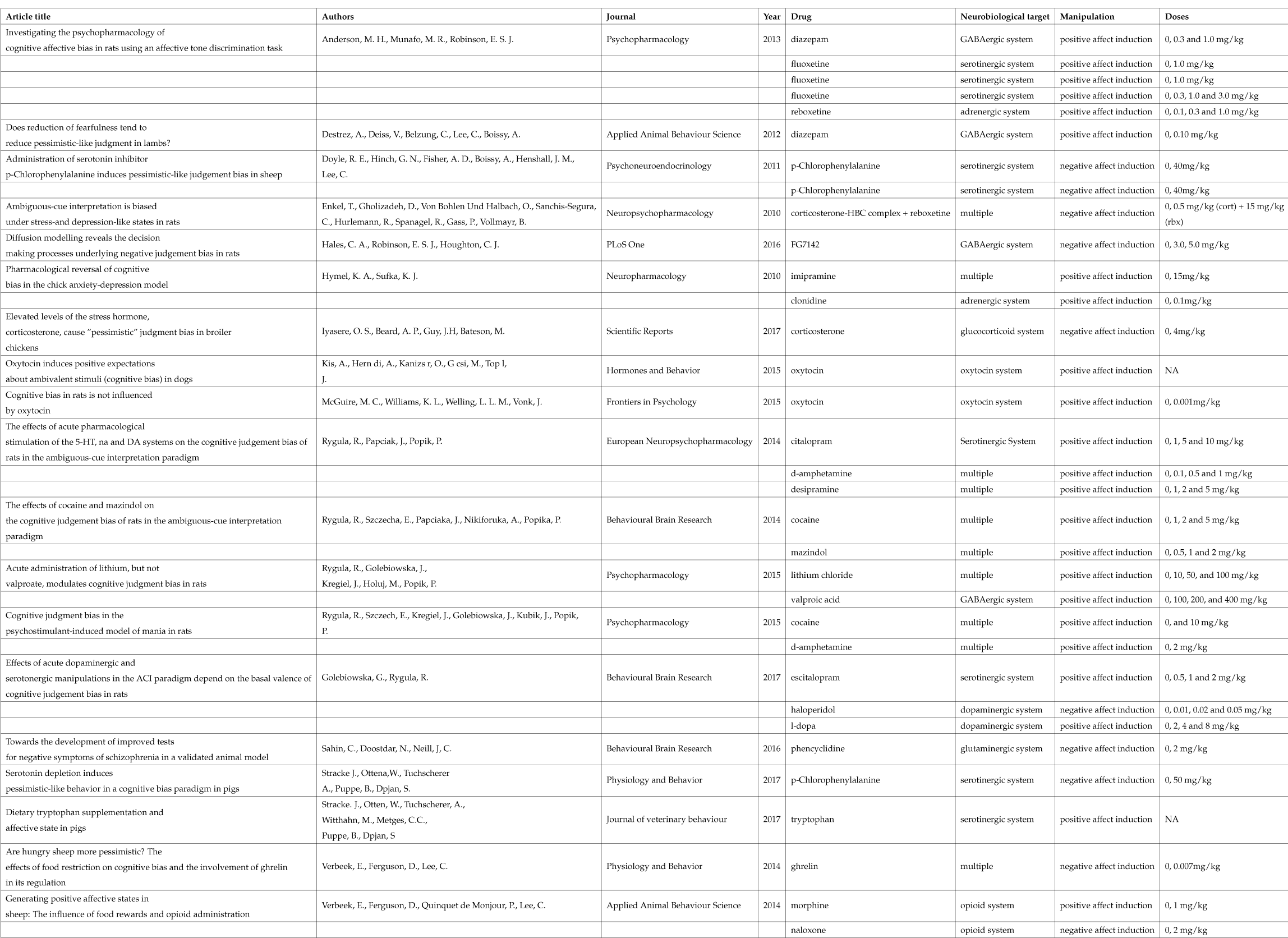
Articles included in the meta-analysis

### 2.4. Effect size and sampling variance calculation

As latency data are bounded at zero, and proportion data are bounded between zero and one, data obtained from the judgement bias task do not follow a Gaussian or normal distribution. The delta method (Taylor approximation) was used to adjust the extracted mean and (sampling) variance (*sd*^2^) prior to calculating the effect size to account for the non-normality of the raw data [45]. For extracted latency data, which were assumed to follow a log-normal distribution, this adjustment was calculated, via the log transformation, as:

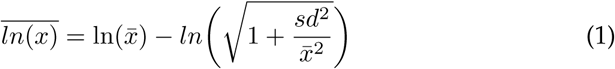

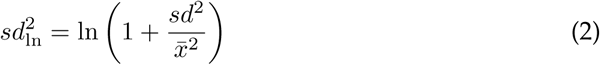

In this case, the transformed sampling variance is exact and not an approximation.

For extracted proportion data, which were assumed to follow a binomial distribution, this adjustment was calculated, via the logit transformation, as [46]:

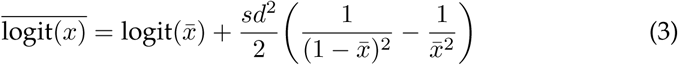

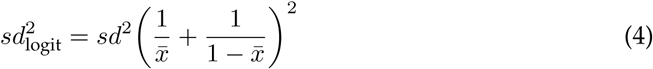

Hedge’s g was then calculated as the difference between the relatively better treatment (in which a relatively more positive affective state was expected) 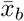 and relatively worse treatment (in which a relatively less positive affective state was expected) 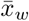 means, divided by the pooled standard deviation, *sd*_*pool*_, and then adjusted for biases arising from small sample sizes by factor *J* which depended on the sample size of the relatively better *n*_*b*_ and relatively worse *n*_*w*_ groups:

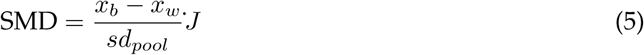

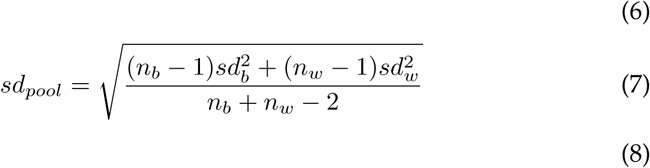

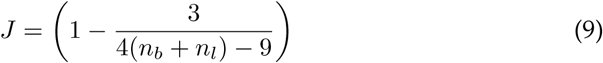

For the latency data, Hedge’s g was multiplied by minus one to account for a higher proportion being equivalent to a lower latency, in terms of judgement bias.

The sampling variance is calculated as follows:

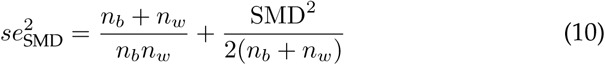

To account for shared controls, if one vehicle treatment group was compared to multiple drug treatment groups, an additional effect size and sampling variance was calculated based on a sample size for the vehicle group that had been divided by the number of treatment groups [47].

### 2.5. Meta-analysis and meta-regression models

The meta-analysis and meta-regression were conducted using the function, rma.mv from the R package metafor [48]; this function allowed us to fit multilevel meta-analytic and meta-regression models [49]. All models included drug, institution at which the research was conducted, and effect ID (a unique ID given to each effect size) as random effects to account for the non-independence of effect sizes from studies conducted at the same institute or using the same drug, and were fit using restricted maximum likelihood. The Knapp and Hartung adjustment was applied to all analyses [50]. To begin with, an intercept only model was fit to the effect sizes. A p-value for this model was obtained using a Wald-type test based on a t-distribution. Heterogeneity was assessed by calculating the *I*^2^ values for each random effect in the model and an overall *I*^2^ value for the model, following [51], which is an extension of the original *I*^2^ [52]

Meta-regression was used to examine whether the following categorical and continuous moderators significantly contributed to variation between effect sizes; the dosing duration (chronic, acute, or chronic wash-out), the time between administration and testing (acute studies only), the number of days since the first dose (chronic studies only), the dosage differences between treatments from which the effect size was calculated, the neurobiological target of the drug, the manipulation type (positive or negative affect induction), the species tested, and the outcome variable used (latency or proportion), presented cue (positive reference cue, near-positive probe cue, midpoint probe cue, near-negative probe cue, negative reference cue), sex of the experimental subjects (all male, all female, or both male and female), reinforcement type (reward-punishment, reward-null, or reward-reward), response type (go/no-go, or go/go), and cue type (reference or probe). An omnibus test based on an *F* distribution was used to assess the significance of each moderator. To further investigate significant moderators, pairwise comparisons were made between the mean effect size for each level of the moderator. A Wald-type test was used to assess the significance of these pairwise comparisons. Moderators which were significant in the meta-regression were subsequently included together in a full model and their influence on the effect sizes was re-assessed. Akaike’s criterion was calculated for the full model and was compared to models where a moderator had been removed, to verify that the model of best fit included all moderators.

### 2.6. Subgroup analyses

As affect is hypothesised to provide information about the probability of each potential outcome of a decision, particularly when there is greater uncertainty about the potential outcomes, any treatment designed to influence affective state is expected to have the greatest influence on judgement bias at the ambiguous cues (see Fig. 2 for example of hypothesised data) [53, 1]. There are methodological and theoretical reasons for why an effect may be observed at one cue and not others. For example, a cue may be too perceptually similar to either of the reference cues for there to be ambiguity about the outcome, or a potential punisher may be much more aversive than the reward is rewarding, to the extent that all animals will avoid probe cues that are similar to the negative reference cue. By considering all cues equally, the effect of an affective manipulation might be obscured, potentially leading to the false inference of no significant effect. To this end, we conducted an additional analysis on a subset of data that included only the effect sizes from the probe cue with the largest absolute effect size for each drug within an article. Additionally, we analysed a second subset of data that included only the effect sizes for the cue with the absolute largest effect size in the direction of the mean effect size for each drug within an article to avoid including outlying effects that might not necessarily reflect the influence of the affective manipulation. If only one probe cue was presented in a study, data from this probe cue were included in the subset data.

**Figure 2:**
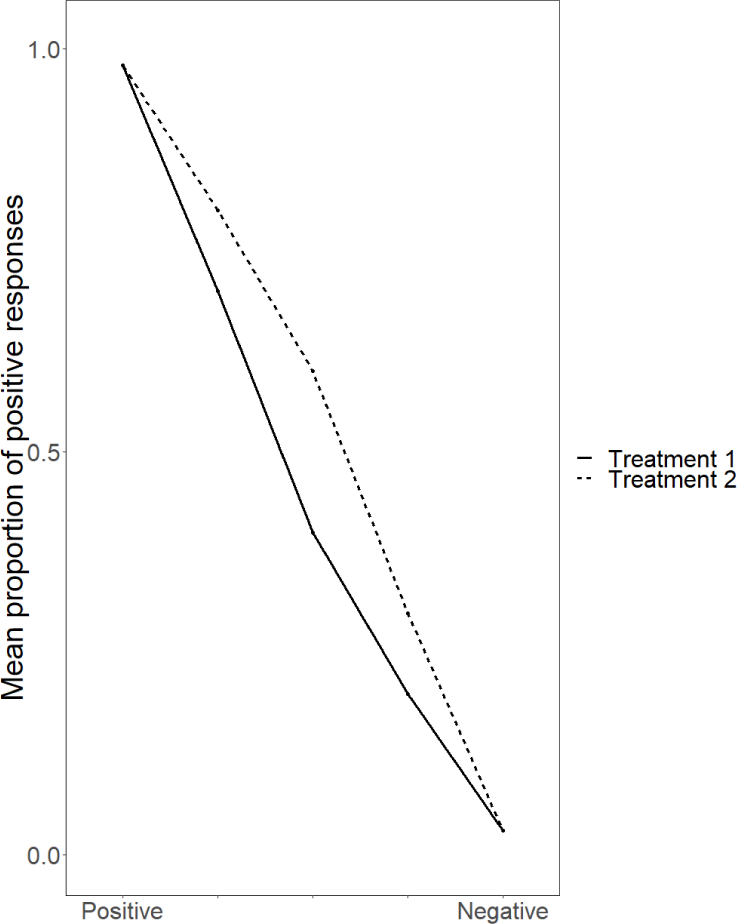
Example of hypothesised data from the judgement bias task with two treatments; one designed to induce a relatively positive affective state (treatment 1) and another designed to induce a relatively negative affective state (treatment 2). While the mean proportion of positive responses is almost identical at the positive and negative reference cue, a treatment difference is observed at the ambiguous cues.

### 2.7. Publication bias and sensitivity analysis

To assess the reliability of results across different analytical approaches and to check for a publication bias, the intercept-only and full meta-regression model were re-fit to the data under a Bayesian statistical framework using the R package MCMCglmm [54]. The non-independence of effect sizes can also be accounted for using Bayesian methods. A parameter-expanded prior was used for both the random effect of drug and institution ID, while the prior variance for random effect of effect ID was fixed at one. Model fitting had 110,000 iterations, 10,000 burn-in periods, and thinning by every 100. The result of this intercept-only model was compared to our initial intercept-only model. The ‘meta-analytic residuals’ (*sensu* [51]) from full meta-regression model conducted in MCMCglmm were used to produce a funnel plot and run Egger’s regression and hence check for a publication bias. Additionally, the intercept-only meta-analysis was repeated but with the effect size and sampling variance that had been adjusted (via the sample size) for shared controls, to assess whether this altered the results.

## 3. Results

### 3.1. Data review

We extracted 440 effect sizes from 19 articles that had been published by authors based at 10 different institutions. Twenty-five different drugs were used across these studies. There were 315 effect sizes (13 articles) that came from studies using acute pharmacological manipulations, 79 effect sizes (5 articles) from studies using chronic pharmacological manipulations, and 46 effect sizes (4 articles) that came from the wash-out period of a chronic pharmacological manipulation (Fig. 3a). Most effect sizes came from studies using drugs that targeted a range of neurobiological systems (164 effect sizes, 7 articles) which included drugs such as cocaine and d-amphetamine which target the dopaminergic, serotoninergic, and adrenal systems, and a large number of drugs specifically targeted the serotonergic system (129 effect sizes, 6 articles) (Fig. 3b). The remaining effect sizes were from experiments using drugs that specifically targeted GABAergic sytem (46 effect sizes, 4 articles), adrenergic system (34 effect size, two articles), dopaminergic system (18 effect sizes, 1 article), opioid system (20 effect sizes, 1 article), glucocorticoid system (15 effect sizes, 1 article), oxytocin system (9 effect sizes, 2 articles), and glutaminergic system (5 effect sizes, 1 articles) (Fig. 3b). The majority (328) of the effect sizes came from studies that had used drugs expected to induce a relatively positive affective state (anxiolytics or antidepressants, 12 articles), while the remainder used anxiogenic or depressant drugs (112 effect sizes, 9 articles). Five different species were used across the studies; the most frequently used species according to the number of effect sizes was rat (301 effect sizes, 10 articles), followed by pig (60 effect sizes, 2 articles), sheep (50 effect sizes, 4 articles), chicken (23 effect sizes, 2 articles), and dog (6 effect sizes, 1 article) (Fig. 3c). Proportion was more commonly used as the outcome measure (318 effect sizes, 11 articles) compared with latency (122 effect sizes, 8 articles). The majority of effect sizes came from studies using only male subjects (304 effect sizes, 10 articles), followed by only female subjects (130 effect sizes, 8 articles), and six effect sizes (1 articles) came from studies that used both male and female subjects (Fig. 3d). The most common reinforcement type was reward-punisher (420 effect sizes, 16 articles) and very few effect sizes came from studies using reward-null (6 effect sizes, 1 article) or reward-reward reinforcement (14 effect sizes, 2 articles) (Fig. 3e). There were more effect sizes from studies using a ‘go/go’ design (298 effect sizes, 9 articles) compared with a ‘go/no-go’ design (142 effect sizes, 10 articles).

**Figure 3:**
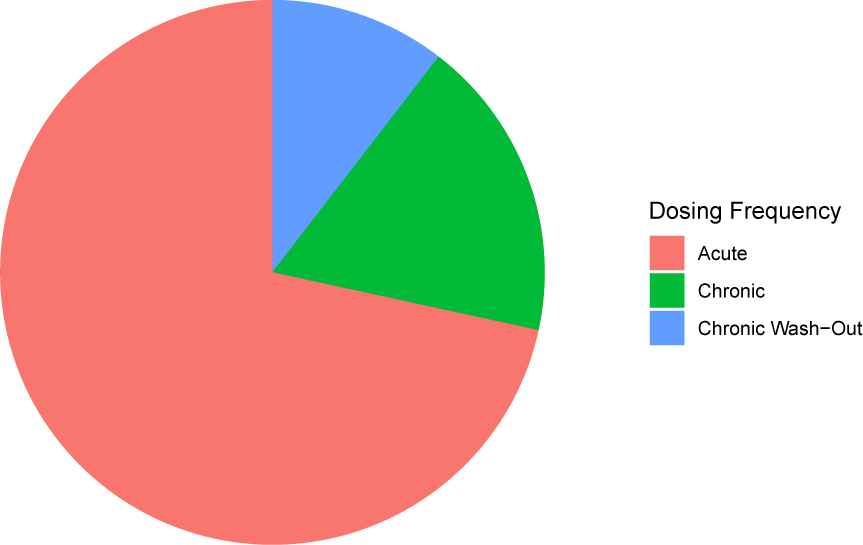
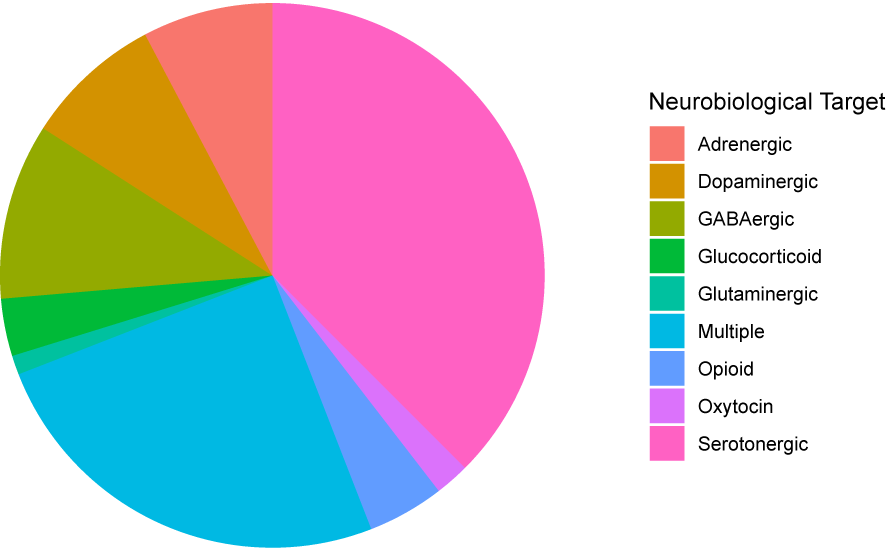
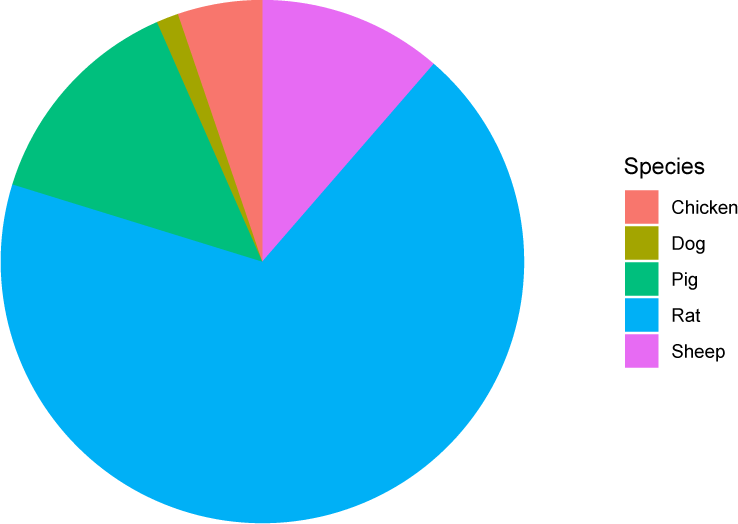
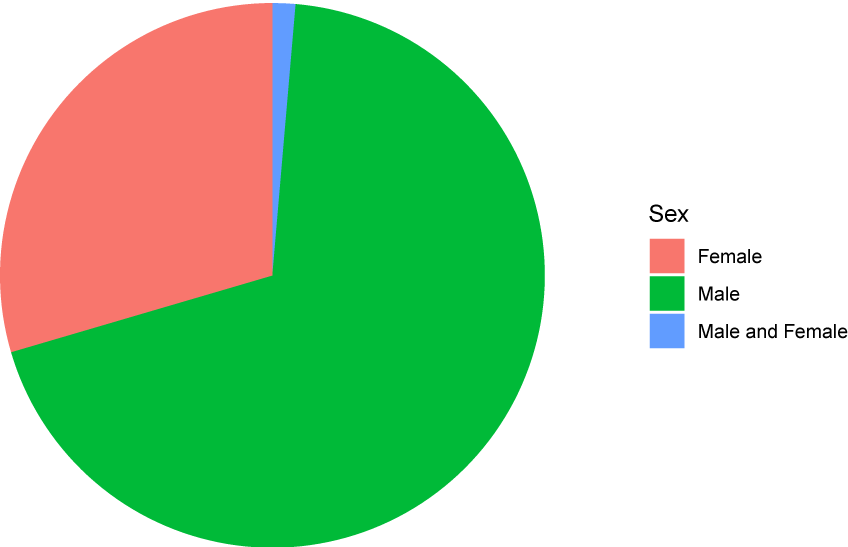
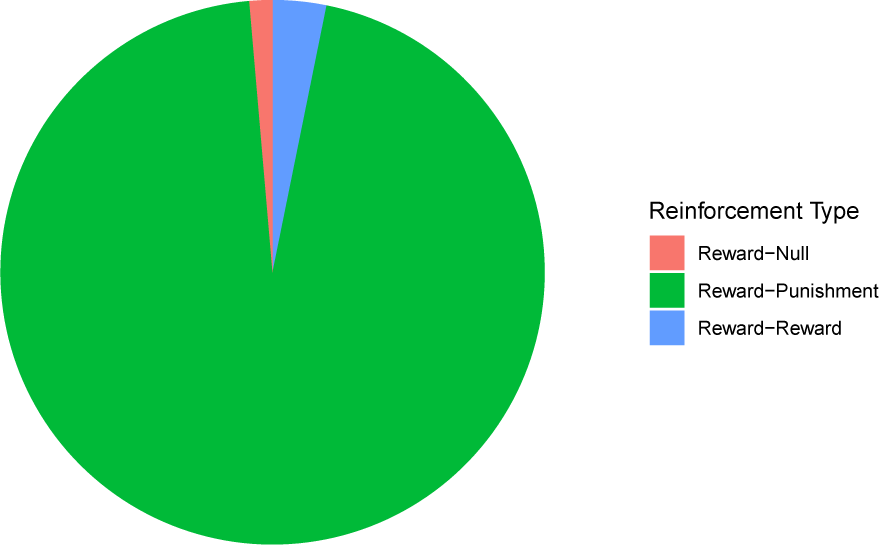
Pie charts illustrating the proportion of effect sizes from each level within a) dosing frequency, b) neurobiological target, c) species, d) sex, and e) reinforcement type.

Across the articles from the acute studies, the average time between the administration of the drug and testing was 38.000*±*2.845 (mean*±*SD) minutes. The average number of days between the start of the chronic drug treatment and testing was 8.556*±* 0.287, and the average days the animal had been withdrawn from a drug when tested in the wash-out period was 7.143*±*0.246. There were 11 articles that used more than one probe cue and three of these articles examined the effect of more than one drug. In total, there were 14 sets of effect sizes obtained from different articles using different drugs which used more than one probe cue. The probe cue with the greatest absolute effect size was the near-positive probe cue on nine occasions, the near-negative probe cue on four occasions, and the midpoint probe cue on one occasion. The probe cue with the greatest absolute effect size was also the presented cue with the greatest absolute effect size in the direction of the mean effect for all but one of the sets of effect sizes, where the near-positive probe cue had the greatest absolute effect sizes and the near-negative probe cue had the greatest absolute effect size in the direction of the mean effect.

### 3.2. Meta-analysis

Overall, drugs expected to alter affective state did not significantly induce a judgement bias (mean=0.127, 95% confidence interval or CI=-0.131-0.384, *t*_439_=0.967, p=0.334). However, this needs to be interpreted in the context of the observed high total heterogeneity in the model, with an *I*^2^ value of 92.854. The between-effect-size effect (i.e., residuals; 44.016%) and the between-drug effect (i.e., which drug were used; 48.839%) explained a large percentage of this heterogeneity, while a very small percentage of variability was due to institutional variation (*<*0.001%). Heterogeneity between effect sizes was further explored through the meta-regression.

However, pharmacological manipulations of affect were found to have a marginally non-significant small to moderate effect (*sensu* [55]: small=0.20, moderate=0.5, large=0.8) on judgement bias when the analysis was repeated on the subset data comprising only data from the ambiguous cue with the largest absolute effect size (mean=0.341, CI=-0.046-0.727, *t*_115_=1.745, p=0.084), and a significant small to moderate effect on the subset data comprising only data from the ambiguous cue with the largest absolute effect size in the direction of the mean effect (mean=0.421, CI=0.011-0.832, *t*_115_=2.032, p=0.045).

### 3.3. Meta-regression

The meta-regression revealed that manipulation type (Fig. 4: *F*_(1,438)_=9.442, p=0.002), the neurobiological target of the drug (Fig. 4: *F*_(8,431)_=2.575, p=0.009), dosage (Fig. 4: *F*_(1,402)_ =10.812, p=0.001), presented cue (Fig. 4: *F*_(4,438)_=2.715, p=0.030), and the cue type (Fig. 4: *F*_(1,438)_=6.655, p=0.010) significantly explained the heterogeneity observed among the extracted effect sizes. Pharmacological manipulations expected to induce a relatively negative affective state (either depressant or anxiogenic) had a greater effect on judgement bias than those expected to induce a relatively positive affective state (Table 2). The effect of drugs targeting the adrenergic system differed significantly from all other drugs used, apart from drugs targeting the oxytocin system for which there was a trend towards significance (Table 2). Drugs targeting the adrenergic system had the opposite effect than expected; a negative judgement bias was induced when a positive judgement bias was hypothesised. Greater differences in dosage between the relatively better and worse treatments were associated with smaller effect sizes. The effect of the pharmacological manipulation was marginally weaker at the positive reference cue compared to the midpoint probe cue, near-negative probe cue, and near-positive probe cue (Table 2). There was no difference in effect size at the positive reference cue compared with the negative reference cue (Table 2). The effect of the pharmacological manipulation was greater at the probe cues compared with the reference cues (Table 1). The remaining moderators tested were not found to significantly explain variation in effect size. These moderators included: species (*F*_(4,435)_=0.947, p=0.437), dosing frequency (*F*_(2,437)_=0.372, p=0.689), time since last dose (*F*_(1,313)_ =0.184, p=0.668), number of days since first treatment (*F*_(1,77)_=1.378, p=0.244), sex (*F*_(2,437)_=0.305, p=0.738), response type (*F*_(1,438)_=1.187, p=0.277), and reinforcement type (*F*_(2,437)_=1.383, p=0.252), and outcome measure (*F*_(1,438)_=0.059, p=0.808).

**Table 2:** Pairwise comparisons of each level of significant moderators from the meta-regression

| Variable | Model | mean Difference | CI Lower Bound | CI Upper Bound | z-val | p-value |
| --- | --- | --- | --- | --- | --- | --- |
| *Manipulation Type* | Negative - Positive Affect Induction | 0.742 | 0.267 | 1.216 | 3.073 | 0.002 |
| *Neurobiological Target (in reference to vehicle or lower doses)* | Serotonergic - Adrenergic | 1.3749 | 0.5349 | 2.2149 | 3.2171 | 0.0014 |
|  | Dopaminergic - Adrenergic | 1.8045 | 0.7928 | 2.8161 | 3.5058 | 0.0005 |
|  | GABAergic - Adrenergic | 1.6843 | 0.8244 | 2.5442 | 3.8497 | 0.0001 |
|  | Glucocorticoid - Adrenergic | 1.9863 | 0.4321 | 3.5404 | 2.512 | 0.0124 |
|  | Multiple - Adrenergic | 1.6467 | 0.7974 | 2.4959 | 3.8111 | 0.0002 |
|  | Glutamnergic - Adrenergic | 2.188 | 0.7179 | 3.658 | 2.9254 | 0.0036 |
|  | Opioid - Adrenergic | 1.2089 | 0.0878 | 2.33 | 2.1194 | 0.0346 |
|  | Oxytocin - Adrenergic | 1.2993 | -0.2707 | 2.8692 | 1.6266 | 0.1045 |
|  | Serotonergic - Multiple | -0.2717 | -0.8011 | 0.2577 | -1.0088 | 0.3136 |
|  | Dopaminergic - Multiple | 0.1578 | -0.5112 | 0.8269 | 0.4636 | 0.6431 |
|  | GABAergic - Multiple | 0.0376 | -0.6277 | 0.703 | 0.1112 | 0.9115 |
|  | Glucocorticoid - Multiple | 0.3396 | -1.0945 | 1.7737 | 0.4654 | 0.6419 |
|  | Glutamnergic - Multiple | 0.5413 | -0.7896 | 1.8722 | 0.7994 | 0.4245 |
|  | Opioid - Multiple | -0.4377 | -1.3304 | 0.455 | -0.9637 | 0.3357 |
|  | Oxytocin - Multiple | -0.3474 | -1.7985 | 1.1038 | -0.4705 | 0.6383 |
|  | Dopaminergic - Serotonergic | 0.4296 | -0.3076 | 1.1667 | 1.1453 | 0.2527 |
|  | GABAergic - Serotonergic | 0.3094 | -0.3518 | 0.9706 | 0.9196 | 0.3583 |
|  | Glucocorticoid - Serotonergic | 0.6113 | -0.8229 | 2.0456 | 0.8377 | 0.4026 |
|  | Glutamnergic - Serotonergic | 0.8131 | -0.4999 | 2.1261 | 1.2171 | 0.2242 |
|  | Opioid - Serotonergic | -0.166 | -1.022 | 0.69 | -0.3812 | 0.7033 |
|  | Oxytocin - Serotonergic | -0.0756 | -1.527 | 1.3757 | -0.1024 | 0.9185 |
|  | GABAergic - Dopaminergic | -0.1202 | -0.9563 | 0.716 | -0.2825 | 0.7777 |
|  | Glucocorticoid - Dopaminergic | 0.1818 | -1.3538 | 1.7173 | 0.2327 | 0.8161 |
|  | Glutamnergic - Dopaminergic | 0.3835 | -1.0556 | 1.8227 | 0.5238 | 0.6007 |
|  | Opioid - Dopaminergic | -0.5955 | -1.6497 | 0.4586 | -1.1104 | 0.2674 |
|  | Oxytocin - Dopaminergic | -0.5052 | -2.0567 | 1.0464 | -0.64 | 0.5225 |
|  | Glucocorticoid - GABAergic | 0.302 | -1.1803 | 1.7842 | 0.4004 | 0.6891 |
|  | Glutamnergic - GABAergic | 0.5037 | -0.8853 | 1.8927 | 0.7128 | 0.4764 |
|  | Opioid - GABAergic | -0.4754 | -1.4779 | 0.5271 | -0.932 | 0.3519 |
|  | Oxytocin - GABAergic | -0.385 | -1.8838 | 1.1138 | -0.5049 | 0.6139 |
|  | Glutamnergic - Glucocorticoid | 0.2017 | -1.6211 | 2.0246 | 0.2175 | 0.8279 |
|  | Opioid - Glucocorticoid | -0.7773 | -2.3618 | 0.8071 | -0.9642 | 0.3355 |
|  | Oxytocin - Glucocorticoid | -0.687 | -2.5738 | 1.1999 | -0.7156 | 0.4746 |
|  | Opioid - Glutamnergic | -0.9791 | -2.4427 | 0.4846 | -1.3147 | 0.1893 |
|  | Oxytocin - Glutamnergic | -0.8887 | -2.7251 | 0.9477 | -0.9512 | 0.342 |
|  | Oxytocin - Opioid | 0.0904 | -1.5096 | 1.6903 | 0.111 | 0.9117 |
| *Presented cue* | Midpoint - Positive | 0.202 | 0.032 | 0.371 | 2.339 | 0.02 |
|  | Negative - Positive | 0.14 | -0.031 | 0.311 | 1.606 | 0.109 |
|  | Near Negative - Positive | 0.296 | 0.03 | 0.562 | 2.188 | 0.029 |
|  | Near Positive - Positive | 0.365 | 0.101 | 0.63 | 2.713 | 0.007 |
|  | Midpoint - Near Positive | -0.164 | -0.428 | 0.101 | -1.216 | 0.225 |
|  | Negative - Near Positive | -0.226 | -0.492 | 0.041 | -1.663 | 0.097 |
|  | Near Negative - Near Positive | -0.069 | -0.378 | 0.239 | -0.441 | 0.66 |
|  | Negative - Midpoint | -0.062 | -0.233 | 0.109 | -0.713 | 0.476 |
|  | Near Negative - Midpoint | 0.095 | -0.172 | 0.361 | 0.698 | 0.485 |
|  | Negative - Near Negative | -0.157 | -0.425 | 0.112 | -1.148 | 0.252 |
| *Cue type* | Reference - Probe | -0.173 | -0.305 | -0.041 | -2.58 | 0.01 |

**Figure 4:**
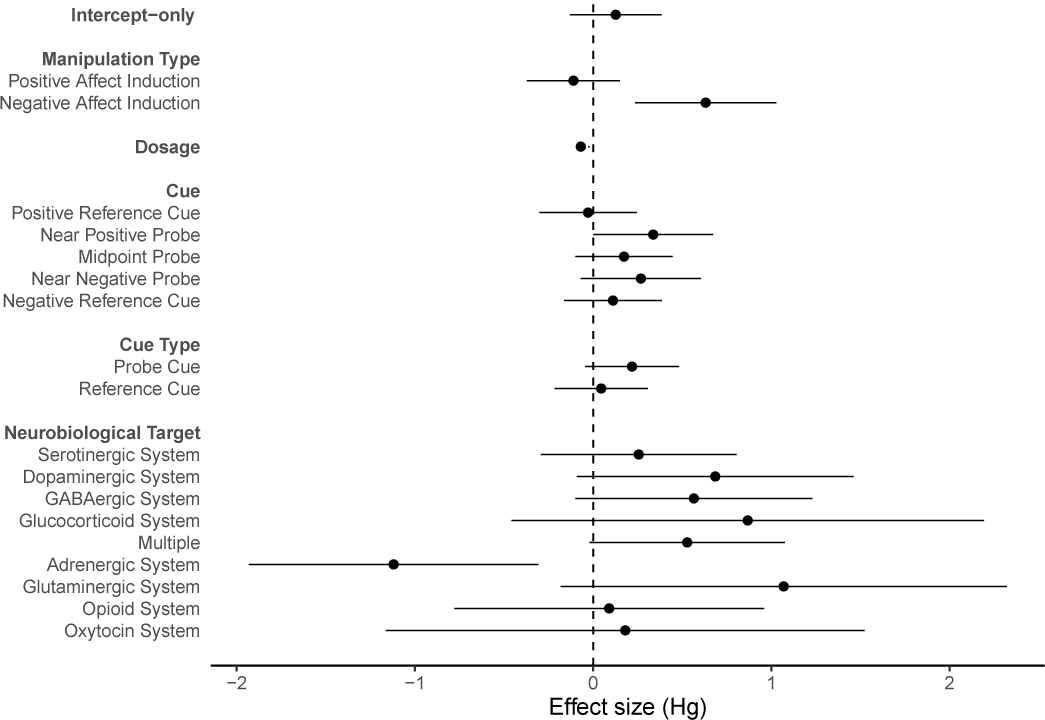
Forest plot with a meta-analytic mean (intercept-only model) and significant moderators from univariate meta-regression models. Each point represents the mean effect size for each moderator and error bars represent the 95% confidence interval.

The best fitting model included cue (i.e. positive reference cue, midpoint probe cue, negative reference cue, near negative probe cue and near positive probe cue) instead of cue type (i.e. reference or probe) (ΔAIC=0.912), and all significant moderators identified in the univariate meta-regression. Removal of the neurobiological target of the drug (ΔAIC=17.746), dosage (ΔAIC=9.837), cue (ΔAIC=4.965), and manipulation type (ΔAIC=3.285) resulted in a poorer fit according to the AIC values. The best fitting model had a marginal *R*^2^ value (*sensu* [56]) of 67.205%. In this model, the difference between effect sizes where a relatively positive compared with relatively negative affective state had been induced was no longer significant, despite having a moderate effect (Δmean= 0.400, CI=-0.165-0.965, t_389_=1.393, p=0.165). Effect sizes from drugs targeting the adrenergic system were overall in the opposite direction to expected and there was a large and significant difference in effect sizes between adrenergic system targeting drugs and serotonergic system targeting drugs (Δmean=1.489, CI=0.628-2.350, *t*_389_=3.400, p*<*0.001), dopaminergic system targeting drugs (Δmean=1.921, CI=0.894-2.948, *t*_389_=3.678, p*<*0.01), GABAergic system targeting drugs (Δmean=1.791, CI=0.917-2.665, *t*_389_= 4.030, p*<*0.001), glucocorticoid system targeting drugs (Δmean=2.041, CI=0.450-3.632, *t*_389_=2.523, p=0.012), multiple system targeting drugs(Δmean=1.734, CI=0.872-2.596, *t*_389_=3.954, p*<*0.001), glutaminergic system targeting drugs (Δmean=2.244, CI=0.734 3.75,3 *t*_389_=2.922, p=0.004), as well as opioid targeting drugs (Δmean=1.279, CI=0.129 2.429, *t*_389_=2.187, p=0.029). There was a large but marginally non-significant difference between the effect sizes of drugs targeting the adrenergic and oxytocin system (Δmean=1.5598, CI=-0.221-3.341, *t*_389_= 1.722, p=0.086). Effect sizes at the positive reference cue were not significantly different from effect sizes at the negative reference cue (Δmean=0.144, CI=-0.037-0.325 *t*_389_=1.563, p=0.119). However, effect sizes were weaker at the positive reference cue compared with the other presented cues. There was a significant and small difference between effect sizes at the positive reference cue and those at the midpoint probe cue (Δmean= 0.200, CI=0.021-0.380, *t*_389_=2.190, p=0.021), and near-positive probe cue (Δmean=0.346, CI=0.052-0.640, *t*_389_=2.312, p=0.021), and a small but marginally non-significant difference at the near-negative probe cue (Δmean=0.271, CI=0.0225-0.567, *t*_389_=1.798, p=0.073).

### 3.4. Study exclusion

As the initial analysis revealed that drugs targeting the adrenergic system had the opposite effect on judgement bias than hypothesised, and there is conflicting evidence about the affect altering properties of adrenergic system targeting drugs in non-human animals [57, 58], we made the post-hoc decision to re-analyse the data excluding adrenergic-system targeting drugs. Two studies had used adrenergic-system targeting drugs, one had used clonidine and the other reboxetine, both of which are considered to induce a relatively positive affective state. These studies accounted for 7.727% (34) of the effect sizes analysed.

### 3.5. Post-exclusion meta-analysis

Following the exclusion of these effect sizes, a small to medium overall effect was observed; pharmacological manipulations of affective state were found to significantly influence judgement bias in the predicted direction (mean=0.433, CI=0.106-0.760, *t*_405_=2.604, p=0.009). However, there again existed high heterogeneity (*I*^2^=91.766%); with 47.708% attributable to between-effect-size effects, 22.404% to between-drug effects, and 21.654% to institutional variation. The meta-analysis using both data subsets (using only one probe cue) revealed a significant and medium overall effect of pharmacological manipulations of affect on judgement bias (absolute greatest probe cue effect sizes: mean=0.526, CI=0.119-0.934, *t*_108_=2.560, p=0.012; and absolute greatest probe cue effect in direction of mean: mean=0.591, CI=0.168-1.014, *t*_108_=2.604, p=0.007).

### 3.6. Post-exclusion meta-regression

While manipulation type (Fig. 5: *F*_(1,404)_=5.982, p=0.015), cue (Fig. 5: *F*_(4,401)_=2.529, p=0.040), and dose (Fig. 5: *F*_(1,368)_=8.647, p=0.004), remained significant as moderators when studies using adrenergic system targeting drugs were excluded, drug target (*F*_(7,398)_=0.569, p=0.782) and cue type (*F*_(1,404)_=2.486, p=0.116) were no longer significant. The model which included all three significant moderators provided a better fit than the models which excluded manipulation type (ΔAIC=8.549), cue (ΔAIC=5.002), and dose (ΔAIC=9.096). This full model had a marginal *R*^2^ value of 57.503%.

**Figure 5:**
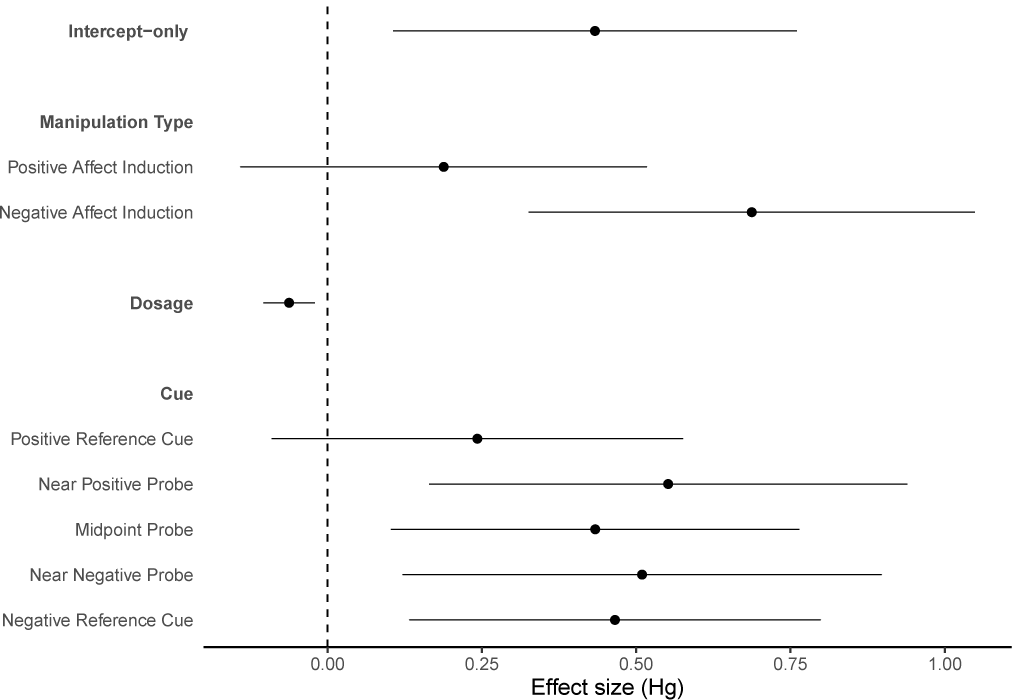
Forest plot with a meta-analytic mean (intercept-only model) and significant moderators from univariate meta-regression models following the exclusion of adrenergic system targeting drugs. Each point represents the mean effect size for each moderator and error bars represent the 95% confidence interval.

Contrary to the previous analysis including adrenergic-targeting drugs, a small and significant difference was found between effect sizes at the negative and positive reference cues, with effect sizes being greater at the negative reference cue (Δmean= 0.236, CI=0.059-0.413, *t*_363_=2.616, p=0.009). The difference in effect sizes between the positive reference and near-negative probe cue was small but no longer significant (Δmean= 0.220, CI=-0.096-0.534, *t*_363_=1.368, p=0.172), which perhaps explains why cue type no longer explained significant variation among effect sizes. The difference between effect sizes at the midpoint probe cue and positive reference cue was very small but significant (Δmean=0.191, CI=0.014-0.367, *t*_363_=2.125, p=0.034), and the difference between effect sizes at the positive reference cue and near-positive probe cue was small but non-significant (Δmean=0.265, CI=-0.050-0.579, *t*_363_=1.657, p=0.099). Effect sizes were still observed to be greater when the anxiogenic or depressant drugs were used compared to the antidepressant or anxiolytic drugs with a moderate difference in effect sizes (Δmean=0.496, CI=0.067-0.926, *t*_363_=2.273, p=0.024), and effect sizes remained significantly greater when there were smaller differences in dosage between the better and worse treatment, although the effect was very small (mean=-0.061, CI=-0.103–0.019, *t*_363_=-2.879, p=0.004).

### 3.7. Publication bias and sensitivity analysis

The results of the Bayesian meta-analysis were consistent with the results of our likelihood-based meta-analyses both prior to and following the removal of effect sizes from studies using drugs targeting the adrenergic system. The effect of the pharmacological manipulations on judgement bias was not significant prior to data exclusion (mean=0.214, 95% credible interval or CI=-0.195-0.586, p=0.236), but a significant overall effect emerged following data exclusion from studies using adrenergic-system targeting drugs (mean=0.440, CI=0.050-0.904, p=0.034).

Visual inspection of the funnel plots produced from the meta-analytic residuals and raw effect sizes (Fig. 6) did not indicate that a publication bias was present, nor did the results of Egger’s test on either the analysis prior to (*t*_402_=- 0.425, p=0.671) or following (*t*_368_=0.384,p=0.701) the exclusion of data.

**Figure 6:**
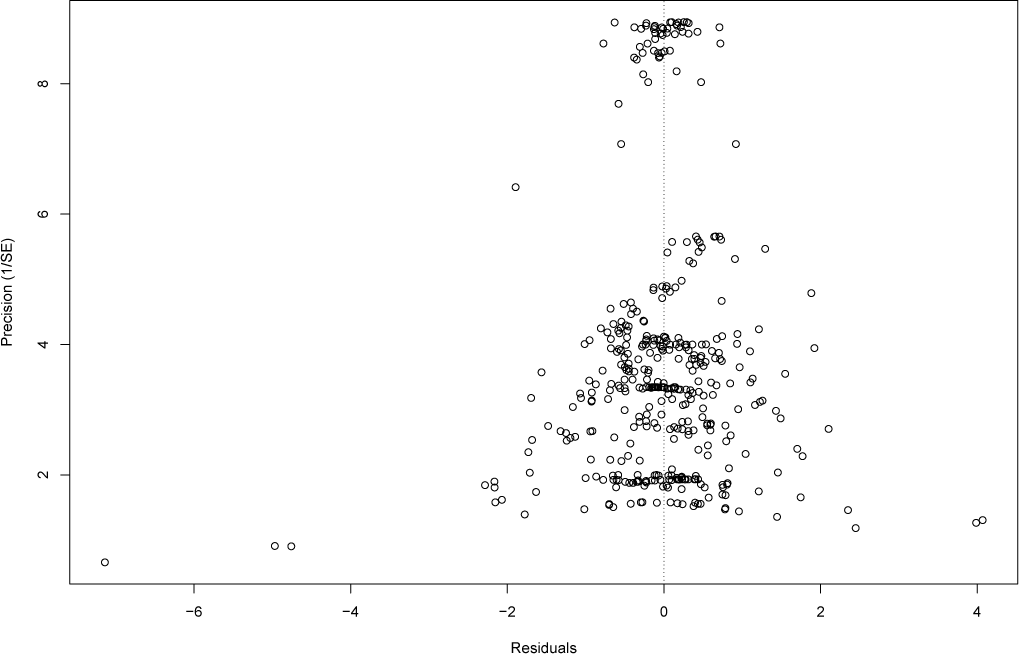
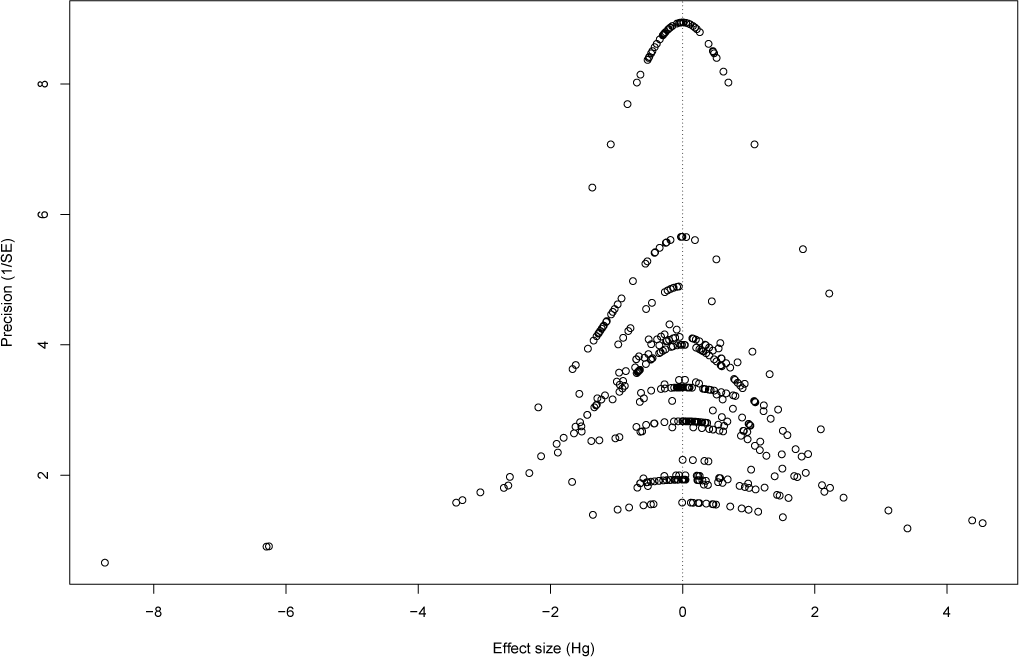
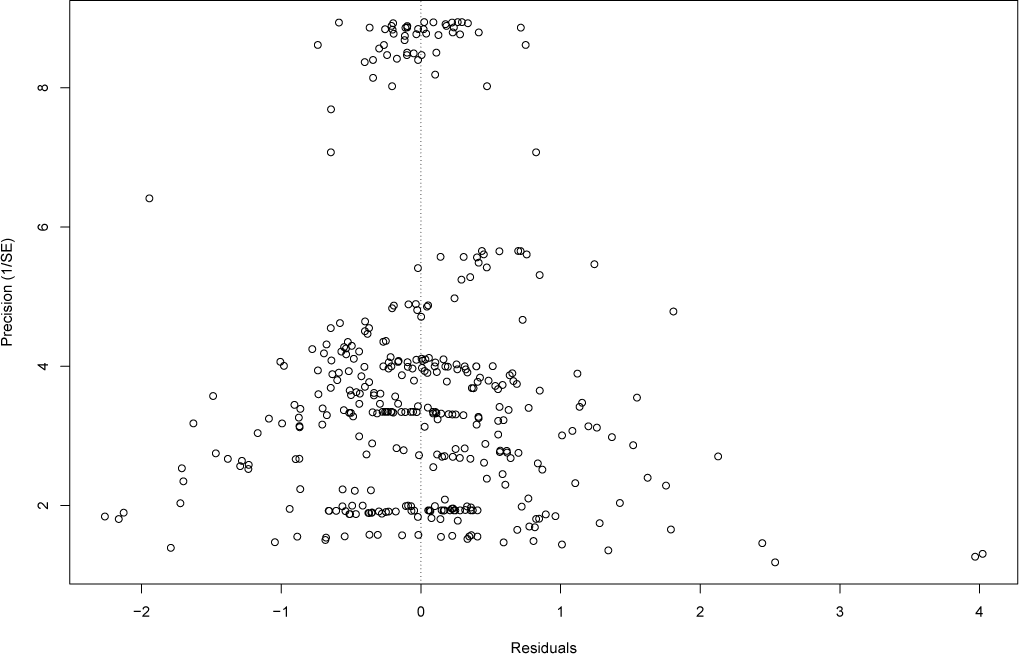
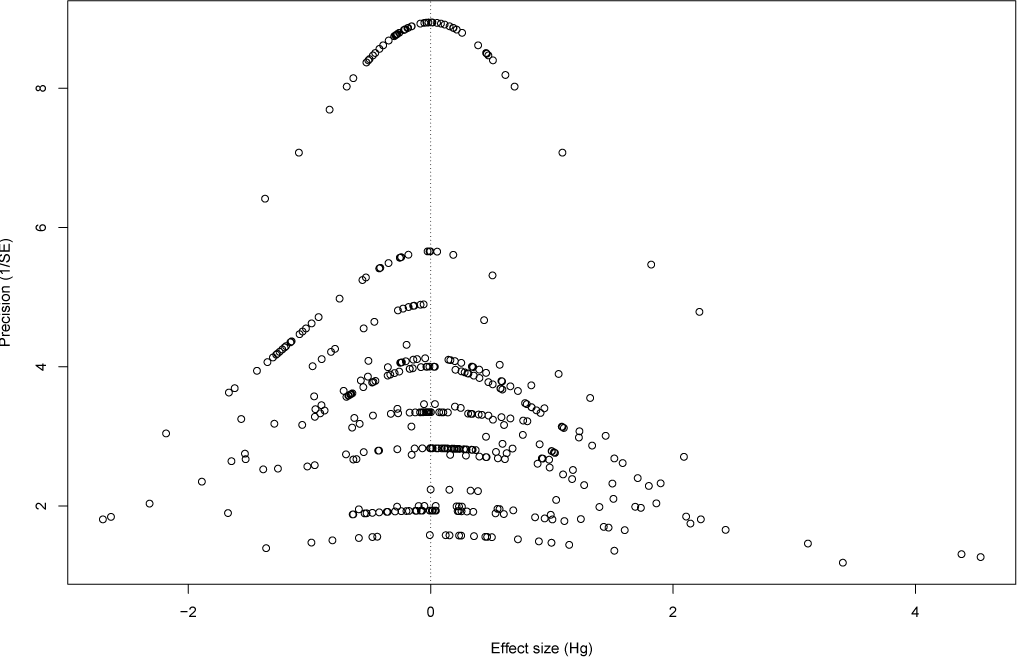
Funnel plots of a) the meta-analytic residual values (residuals + sampling errors) for the full meta-regression model prior to exclusion of effect sizes from studies using adrenergic system targeting drugs; b) the raw effect sizes and the inverse standard errors prior to exclusion of effect sizes from studies using adrenergic system targeting drugs; c) the meta-analytic residual values for the full meta-regression model following exclusion of effect sizes from studies using adrenergic system targeting drugs; d) the raw effect sizes and the inverse standard errors following exclusion of effect sizes from studies using adrenergic system targeting drugs.

Re-analysis of the intercept-only model using the effect sizes and variances that had been adjusted for shared controls did not alter the results qualitatively. The result prior to data exclusion was statistically non-significant (mean=0.127, CI=-0.131-0.385, *t*_439_=0.965,p=0.335) while following data exclusion was significant (mean=0.435, CI=0.129-0.741, *t*_410_=2.79, p=0.006).

## 4. Discussion

Judgement bias is a relatively new and promising measure of animal affect that may provide a useful alternative to more common behavioural assays used to assess the efficacy of potential pharmacological treatments of mood disorders, such as the forced swim test. Empirical studies with human subjects have supported its construct validity [26, 27, 28, 29, 30]. To examine its predictive validity, we conducted a systematic review and meta-analysis of studies investigating the effect of affect-altering drugs on judgement bias in non-human animals. We analysed data from 19 published research articles which yielded 440 effect sizes.

There was high heterogeneity (>90%) between the effect sizes observed [59] indicating strong variability in the extent to which pharmacological manipulations of affective state alter judgement bias. The drug used accounted for some of this variability, as did the institution at which the research was conducted, yet a high proportion of heterogeneity was also attributed to variation within drug and institution. Our meta-regression further highlighted a number of factors which explained variation in effect sizes including the neurobiological drug target, manipulation type (whether the drug was hypothesised to induce a negative or positive affective state), dosage, cue, and cue type (reference or probe).

Although our initial overall meta-analytic model found no significant overall effect of affect-altering drugs on judgement bias, the results of the meta-regression showed a clear moderating effect of the neurobiological drug target, particularly of drugs targeting the adrenergic system. A small to medium effect was found following the removal of data from studies targeting the adrenergic system. These two excluded studies used reboxetine, an antidepressant, and clonidine, which is used off-licence to treat anxiety disorders. Jointly, these drugs were found to exert an opposite effect on judgement bias; inducing a negative judgement bias when a positive judgement bias was predicted. Paradoxically, depression and anxiety are known side effects of these drugs [60]. Moreover, these studies both used an acute dose. Both norepinephrine and cortisol increase in response to stress and acute dosing of drugs which simultaneously elevate levels of norepineprine and cortisol have been shown to result in stress-like changes in the neural response to negative stimuli in humans [61]. It is therefore feasible that the acute delivery of adrenergic-system targeting antidepressant drugs induced a relatively negative rather than a positive affective state which resulted in the relatively negative judgement bias observed. This potential explanation is further supported by studies that have observed anxiety-like states in rodents following the administration of similar adrenergic-system targeting drugs [57, 58].

An alternative explanation could be related to another side effect of adrenergic agonists documented in human and non-human animal subjects; sedation [62, 60] A sedated animal may not have been able to fully partake in the experiment or have been considerably slower to respond, leading to seemingly ‘pessimistic’ responses. However, further studies should be conducted to reveal the extent to which the results from these two studies can be generalised to all adrenergic system targeting drugs.

Drugs inducing negative affect (e.g. depressants and anxiogenics), had a greater effect on measured judgement bias than drugs inducing positive affect (e.g. antidepressants and anxiolytics). This result may reflect an interaction between the drugs administered and affective states arising from the process of being tested, which may sometimes be negative in their own right (e.g. invasive administration of drugs, social isolation during testing). These factors may have enhanced the effect of the negative-affect inducing drug, while dampening the effect of the positive-affect inducing drug. Indeed, in humans, there is evidence to suggest that some affect-altering recreational drugs intensify the affective state of an individual prior to consumption, or result in the exaggerated interpretation of emotional stimuli [63, 64].

Another possible explanation for the moderating effect of manipulation type is that there are floor effects which limit the impact positive-affect inducing drugs can have on judgement bias. There will be a physical limit to how quickly an animal can approach a cue, and the control animals may already be performing at or close to this limit, meaning that the animals that had been given the positive drug could not respond any quicker. However, this explanation will only be relevant to studies measuring approach latency. Similarly, the smoke-detector principle states that individuals should be overly responsive to potential threats [65]. An individual may continue to err on the side of caution even when in a more positive affective state as avoiding punishers is likely to be perceived as less costly than incurring them [65, 66].

Greater effects were observed when there were relatively smaller differences in dosage between treatments. This is consistent with the inverted U-shaped dose-response function that is sometimes observed in drug studies [67, 68, 69]. This result may reflect that higher doses increase the probability of side effects which may interfere with task performance [70, 71]. Adverse effects that alter the motivation of the animal (e.g. reduced appetite), their consummatory behaviour (e.g. nausea), or psychomotor abilities (e.g. sedation) are likely to affect judgement bias. Such side effects are common to several affect-altering drugs [60]. It may therefore be sensible to take measures of activity or food consumption concurrent to the judgement bias task to assess the impact of potential side effects of drug manipulations.

The meta-analysis also found that the effects of the pharmacological manipulations on judgement bias were weakest at the positive reference cue, and the effect of pharmacological manipulations was greater when only the probe cue with the greatest effect size within each drug and article were analysed. This reflects that pharmacological manipulations of affect exert a stronger influence on trials where there is ambiguity about the outcome of the trial compared with trials where the reward is certain. On presentation of the positive reference cue, there should be little ambiguity about the outcome, and it would be expected that the animal should make the response that allows them to obtain the reward on a high proportion of trials. The influence of any affective manipulation should be greatest when there is uncertainty about the outcome as subjective probabilities of uncertain outcomes are thought to be more strongly informed by an individual’s affective state [1, 72, 73]. Thus, this finding is consistent with the theoretical framework underlying judgement bias.

However, it is unclear why the extracted effect sizes were not smaller at the negative reference cue following exclusion of effect sizes from studies that had used adrenergic-targeting drugs. The pharmacological manipulations were not expected to influence responses to the negative reference cue as there should be no uncertainty about the outcome. Moreover, the pharmacological manipulations rarely exerted the greatest influence at midpoint cue, where there should be the greatest uncertainty about the outcome, in studies where multiple probe cues were presented. This further suggests pharmacologically induced affective states do not necessarily induce a greater judgement bias as uncertainty about the decision outcome increases. Both valuation and probability of decision outcomes are key components of decision-making; an individual might be more likely to make a risky or more ‘optimistic’ response either if they considered the reward to more probable or punisher to be less probable, or if they considered the reward to be more valuable or punisher to be less aversive [53, 74]. Speculatively, it is possible that the pharmacological manipulations altered the valuation of the punisher, hence altering responses to its presentation.

Our meta-analysis found no evidence to indicate that the species used, the dosing frequency, the time since last dose in acute studies, the number of days since first treatment in chronic studies, the outcome variable used, the biological sex of the individuals studied, the reinforcement type, or response type had moderating effects on the influence of pharmacological manipulations of affect on judgement bias. While this might reflect that there is insufficient power to detect an effect, it might indicate that judgement bias is robust to variation in methodology and across species. Interestingly, despite being one of the most commonly used non-human animal species in research, none of the studies included in this meta-analysis used mice [75, 76]. We found no evidence to suggest a publication bias.

Future studies should attempt to account for the potential side effects of pharmacological manipulations. Observing behaviour following drug administration, for example activity levels and food and water consumption, may help to highlight potential adverse effects. The majority of effect sizes extracted in this meta-analysis were from studies using serotonergic-system targeting drugs. While this is unsurprising given that commonly prescribed antidepressants target the serotonin system [42], mood disorders are associated with dysfunction in several neurological systems and further investigation of the influence of pharmacologically-induced changes in the activity of these systems may be beneficial [77, 78, 79]. This meta-analysis has highlighted that multiple probe cues may be preferable in future studies. Pharmacological manipulations of affective state do not necessarily exert the strongest influence of judgement bias at the most ambiguous cue, as found in this meta-analysis, and using multiple cues would allow a more comprehensive assessment of the effect of the manipulation. Finally, it would be worthwhile to assess the efficacy of judgement bias as a measure of pharmacological manipulations of affect in mice.

To conclude, this meta-analysis has provided evidence that judgement bias has predictive validity as a measure of the affective impact of pharmacological manipulations. However, a key issue is the potential interference of drug side effects on judgement bias. In particular, the contrary effect of adrenergic-targeting affect-altering drugs and the greater effect of drugs on judgement bias at lower doses, may be attributed to side effects, or to the complex nature of adrenergic drug effects. The effect of drugs hypothesised to induce a negative affective state was greater than the effect of drugs hypothesised to induce a positive affective state, thus larger sample sizes may be required when testing the efficacy of potential pharmacological treatments for mood disorders. However, if consideration is given to these potential shortcomings, the judgement bias task for which there is evidence of construct validity and now of predictive validity, appears to be a viable measure of pharmacologically-induced affect in non-human animals.

## Acknowledgments

We thank the Biotechnology and Biological Sciences Research Council (BB-SRC: SWBio Doctoral Training Programme grant number BB/M009122/1) for funding this work. SN and ML were supported by an Australian Research Council (ARC) Discovery grant (DP180100818).

